# Cognition through internal models: Mirror neurons as one manifestation of a broader mechanism

**DOI:** 10.1101/2022.09.06.506071

**Authors:** Tyson Aflalo, Srinivas Chivukula, Carey Zhang, Emily R. Rosario, Nader Pouratian, Richard A. Andersen

**Affiliations:** Department of Biology and Biological Engineering, California Institute of Technology, Pasadena CA 91125, United States; Tianqiao and Chrissy Chen Brain-Machine Interface Center, Chen Institute for Neuroscience, California Institute of Technology, Pasadena CA 91125, United States; UT Southwestern Medical Center, Dallas, Texas, 75390, United States; Casa Colina Hospital and Centers for Healthcare, Pomona CA 91767, United States

## Abstract

Cognition relies on transforming sensory inputs into a more generalizable understanding. Mirror neurons are proposed to underlie this process, yet they fail to explain many key features of human thinking and learning. Here we hypothesize that mirror-like responses are one limited view into a more general framework by which internal models of the world are built and used. We recorded populations of single neurons in the human posterior parietal cortex as a participant felt or observed diverse tactile stimuli. We found that mirror-like responses were fragile and embedded within a richer population response that encoded generalizable and compositional features of the stimuli. We speculate that populations of neurons support versatile understanding, not through mirroring, but instead by encoding representational building blocks of cognition.

**One-Sentence Summary:** Similar neural responses during observed and experienced sensations are mediated by shared compositional building blocks, not mirror neurons.

## Main Text

We don’t just see the world. We understand it (*1–3*). From a brief video or even a still image of a person in action, we can infer what they are doing, why they are doing it, what they will do next, or what they might have done but didn’t. A fundamental question in neuroscience is how neural populations transform sensory inputs into such deep and versatile understanding.

Mirror neurons have been proposed to be the neural basis for such understanding, at least for how we understand what another person intends or feels (*2, 4*). In this view, we map the visual representation of others’ actions, emotions, or sensations, onto our own corresponding neurons and thereby attain understanding (*2, 4, 5*). However, this hypothesis has received numerous critiques (*6, 7*). For example, if understanding comes from activating our own high-level action representations, how can we understand actions we have never performed (e.g., jumping a skateboard)? Or could never perform (e.g., flying)? Alternatively, a potential role for these neurons may be clouded by an emphasis on interpreting the behavior of single neurons as opposed to neural populations (*8, 9*).

Alternate theoretical frameworks have emerged in parallel based on the concept that human-like learning and thinking is the product of how neural systems build and use internal models of the world, what we will call the “cognition through internal models” framework (*3, 10*). Internal models are our brain’s representations of ourselves and our physical world: the objects in it, how they interact, and how they do not. The power of these internal models lies in defining features, such as generalizability and compositionality. Generalizability captures the idea that representations apply across contexts and behaviors, providing a common substrate to inform our perception, cognition, imagination, and planning across many situations. Compositionality captures the idea that multi-faceted representations are constructed as a combination of more basic level representations (*11, 12*). Recent studies have demonstrated compositional encoding at the neural population level (*13–17*) suggesting that a similar scheme might underly neural responses typically associated with mirror neurons. Such an encoding architecture is highly versatile and better positioned to overcome many of the limitations associated with the mirror hypothesis.

We have recently recorded populations of single neurons in the human posterior parietal cortex (PPC) during motor, sensory, and cognitive behaviors. These neurons encode many diverse body-related variables such as action verbs, observed actions, motor and sensory imagery, and motor plans (*18–21*). Individual neurons are often complex, yet population-level representations demonstrate generalizable encoding across these varied domains in a functional organization we termed partially-mixed selectivity (*20*). Based on these past results, we hypothesize that “mirroring” is one view into a more general mechanism by which we create generalizable internal representations of the world. To test this hypothesis, we recorded from populations of neurons in human PPC while a participant experienced actual touch (to the participant) or observed touch (to another individual). We found that when using rich multi-dimensional stimuli, individual neurons were not well characterized by mirroring. Instead, at the neural population level, basic-level tactile variables related to body location and touch type were encoded as generalizable compositional building blocks embedded within latent neural subspaces. Recent work in the cognitive neuroscience literature hypothesizes that human learning and thinking are largely enabled by how we build and utilize models of the world to understand, explain, imagine, and plan. Human PPC’s neural population responses support this hypothesis by demonstrating that language, imagination, planning, and perception tap into the same underlying neural substrates.

## RESULTS

We recorded populations of single neurons from a microelectrode array implanted in participant NS, a tetraplegic individual (spinal injury at cervical levels 3–4; C3/4) participating in a clinical brain-machine interface study (Fig S1). NS had well-defined sensory receptive fields that responded at short (~60ms) latencies suggesting a role in the bottom-up sensory processing (*19*). Such responses opened the possibility of studying mirror-like phenomena in the sensory domain. We performed two primary experiments: the first used a simple paradigm to confirm mirror-like properties; the second expanded the number of task dimensions to test whether PPC populations support a more general mechanism.

### Mirror-like responses in human PPC single neurons

We found many compelling examples of neurons with mirror-like responses. Supplementary Video 1 ([link here](https://www.youtube.com/watch?v=tOjJHe1eums)) shows an example neuron demonstrating specificity and congruency, the defining features of mirror responses. Specificity is the selective activation to a restricted set of tactile stimuli, evidenced by the neuron responding to touch to her outer shoulder but not her inner shoulder. Congruency is defined as having similar neural responses when experiencing or when observing tactile stimulation. Supplementary video 2 ([link here](https://www.youtube.com/watch?v=39yC-lwbp-s)) shows an additional example.

We performed a basic sensory mirroring task (Experiment 1 – BSMT, Figure 1A) to quantify the existence of sensory mirror-like responses. We recorded an average of 126 ± 20 neurons over 6 sessions while the participant felt rubbing motions applied to her cheek or shoulder or observed rubbing motions applied to an experimenter’s cheek or shoulder. This two-factor (body part x person) design allowed us to test for specificity and congruency. We found robust coding of experienced and observed tactile sensations (Figure 1B, 1C). Many neurons demonstrated mirror-like responses, firing similarly to touches to the cheek or shoulder (specificity), invariant to whether the touches were felt or observed (congruency) (model analysis, body part specific, p<0.05 corrected, Figure 1D). Example neurons showing mirror-like responses are shown in Figure 1E. We summarized the response of the entire population during each condition as a vector of the mean firing rates while the participant experienced or observed touch. We found that population correlations between the observation and experience conditions was higher for matching body parts than for mismatched body parts (t-test, p<0.05, Figure 1F). Thus, the mirroring phenomena was robust at the neural population level.

**Figure 1.**
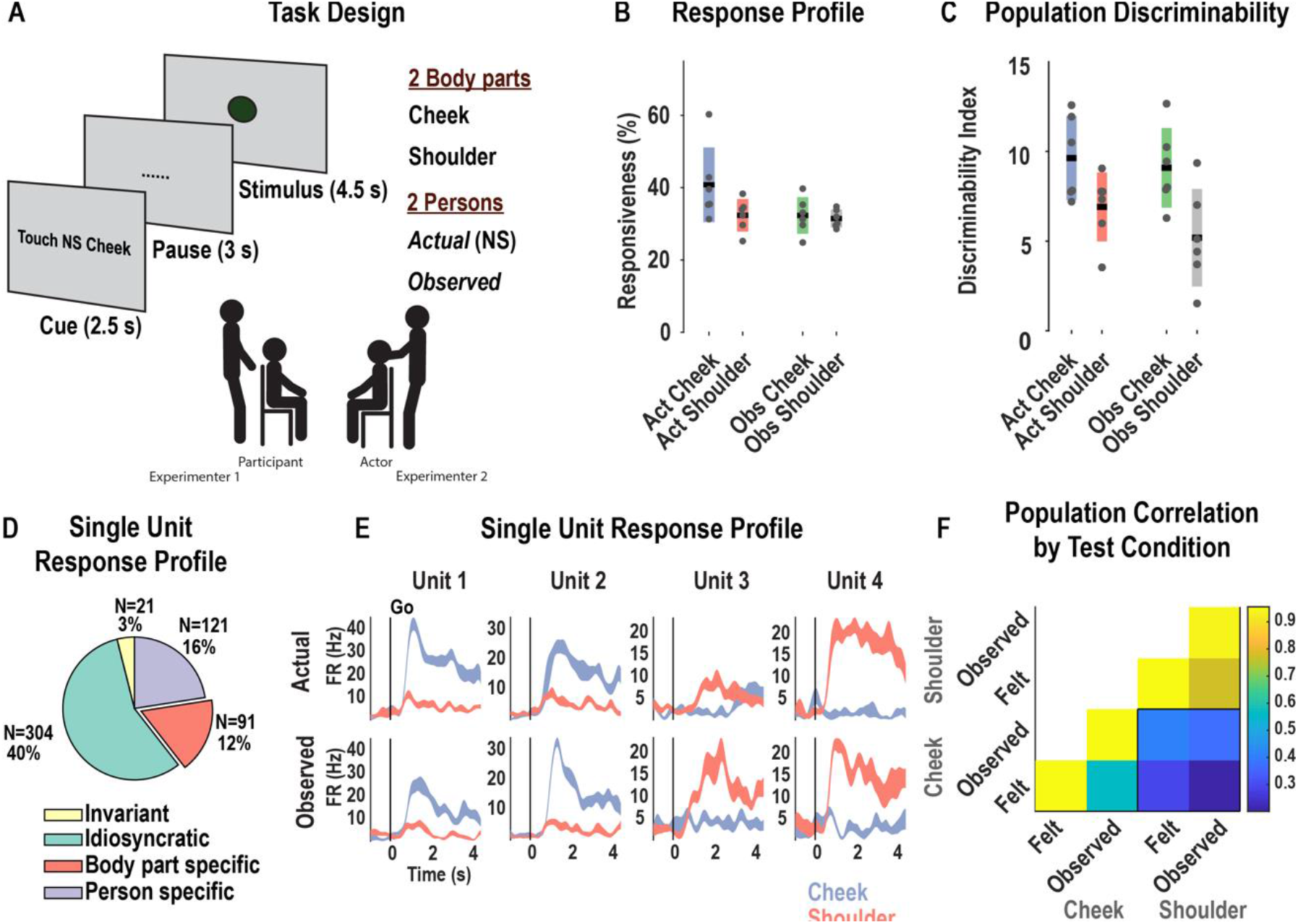
Evidence for mirror-like responses for *actual* and *observed* touch in human PPC. **A**, Task design testing neural responses during experienced and observed touch to the cheek and shoulder. The cue (hidden from the participant) instructed experimenters on which tactile stimulus to deliver during the stimulus phase. **B**, Percent of neurons demonstrating significant modulation from the inter-trial-interval baseline (*p*<0.05, FDR corrected, mean ± 95% CI, 10 trials per condition, 757 neurons). Gray dots represent single session results. Act = actual; obs = observed. **C**, Population measure of the strength of representation as measured by the distance of neural population response from the baseline ITI period baseline (Mahalanobis distance, mean ± 95% CI across sessions). Gray dots represent single session results. **D**, Pie chart categorizing neurons according to their individual response properties: body part specific (invariant to whether touch was actual or observed, i.e. mirror-like), person-specific (invariant to body part), invariant (responsive to all conditions), or idiosyncratic (other patterns). **E**, Example neurons showing mirror-like responses (mean±SEM, n=10 trials.) Each column shows the response for one neuron to actual (top row) and observed touch (bottom row.) **F**, Cross-validated correlation of population responses within and between conditions. Colors represent the correlation strength, as in the scale.

These data provide powerful support that individual neurons can respond in similar ways to experienced and observed sensations. However, from this simple paradigm, it is unclear whether similar responses to actual and observed sensations are direct evidence for a mirror mechanism, or instead, part of a more general mechanism of cognition. Also, already it can be seen that only 12 percent of neurons demonstrate specificity and congruency, indicating that a large proportion do not fit the criteria for mirroring responses.

#### Unpacking the population code: multidimensional sensory mirroring task

To better understand similar encoding between experienced and observed actions, we performed a second experiment (Experiment 2) that augmented the first experiment to include four different types of touch (pinch, press, rub, and tap). These touch-types were selected as they resulted in perceptually distinct stimuli under observed and actual touch conditions and not based on assumptions about the underlying selectivity of recorded neurons. Thus, in the updated task, three manipulated dimensions (body part, touch type, and person) are combined in a full factorial design for 16 total conditions. Including the additional dimension allowed us to 1) test whether encoding similar variables in similar ways was a ubiquitous property of the neural population; and 2) test whether these responses are consistent with a compositional basis, encoding multidimensional sensations as a combination of basic sensory properties. We recorded an average of 119 ± 16 neurons over 8 sessions.

#### Single neuron responses are heterogeneous

As in the first experiment, we found robust coding of experienced and observed tactile sensations (Figure 2A,2B). The response to the different touch types could be discriminated for experienced or observed conditions (time-resolved classification, Figure 2C). However, the inclusion of additional touch-types highlighted the near-universal complexity of single-unit responses: Neurons that appeared to have a simple mirror response for a single touch-type were no longer easily reconciled with a mirror neuron account: Figure 2D shows a neuron that responds similarly to experienced or observed pinches to the cheek, but not the shoulder, consistent with a mirror account. However, testing the same neuron with additional touch-types reveals a more complicated pattern (Figure 2E): the neuron is selective for pinches to her own cheek but responds to all touch-types during observation. A straightforward interpretation of the mirror mechanism would predict that NS would understand all touch-types as a pinch, inconsistent with behavioral evidence that the touch types were easily discriminated and the finding that observed touch types are discriminable (e.g., Figure 2C). Additional example neurons illustrating heterogeneous and complex responses are shown in Figure 2E and Figure S2.

**2.**
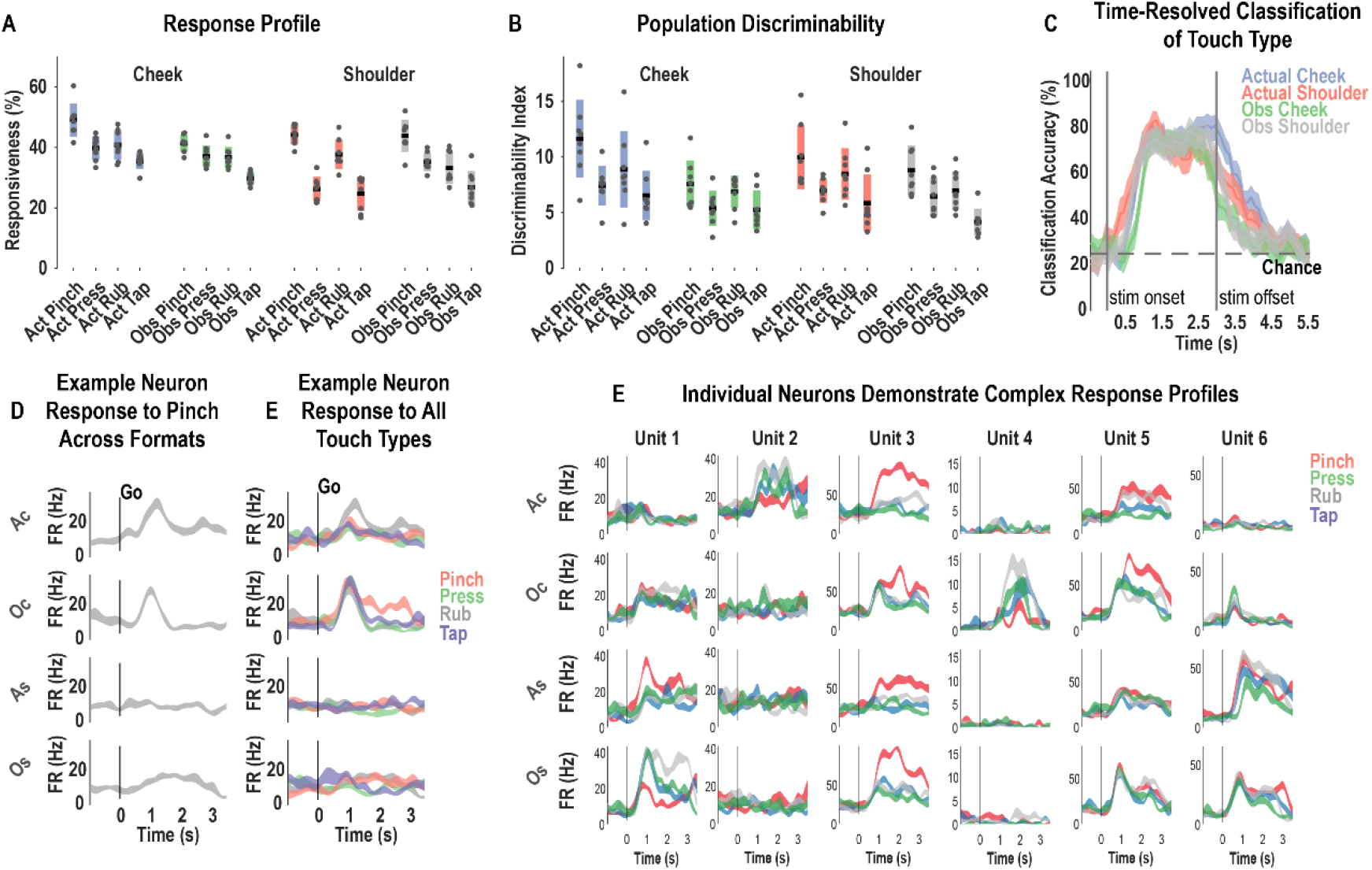
Single neurons discriminate many types of *actual* and *observed* touch. **A**, Percent of neurons demonstrating significant modulation from the inter-trial-interval baseline (*p*<0.05, FDR corrected, mean ± 95% CI, 10 trials per condition, 757 neurons). Gray dots represent single session results. **B**, Population measure of the strength of representation as measured by the distance of neural population response from the baseline ITI period baseline (Mahalanobis distance, mean ± 95% CI across sessions). Gray dots represent single session results. C,Time-resolved, cross-validated classification accuracy discriminating the four touch-types within each sensory field (mean ± 95% CI computed across sessions). **D**, Sample neuron response to pinch across all sensory fields as a function of time (mean ± SEM, n=10 trials.) **E**, Response for the same example neuron from panel D across the four sensory fields, now including all touch types. Touch types are color-coded, as indicated. Other details are as in panel D. **F**, Additional example neurons (see also Figure S2). Each column depicts the response for one unit to each sensory field (rows). Details as in panels D. Act, ***actual***; Obs, ***observed***; s, seconds; Ac, ***actual*** cheek; Oc, ***observed*** cheek; As, ***actual*** shoulder; Os, ***observed*** shoulder; Hz, hertz

We used a model selection analysis (*21*) to categorize patterns of congruency across all sensory fields (e.g., the cheek or shoulder, on NS or the experimenter). For each neuron, we fit linear tuning models that described the response of the neuron to the four touch-types (selectivity pattern, SP) as either congruent or incongruent across sensory fields. There are 51 such possible models (Figure S3). Three schematic examples illustrating congruency patterns are shown in Figure 3A-C. From among the 51 possibilities, we identified the linear model that best described neural behavior using two metrics: Bayesian information criterion (BIC) and cross-validated coefficient of determination (cvR^2^). The percentage of PPC cells that behaved according to each model are shown in Figure S4. Figure 3D summarizes the result by grouping responses into eight general categories that captured high-level modes of behavior (see Figure S5 for the results split by BIC or cvR^2^ criteria). A summary description of the eight modes can be found in the legend of Figure S3.

**Figure 3.**
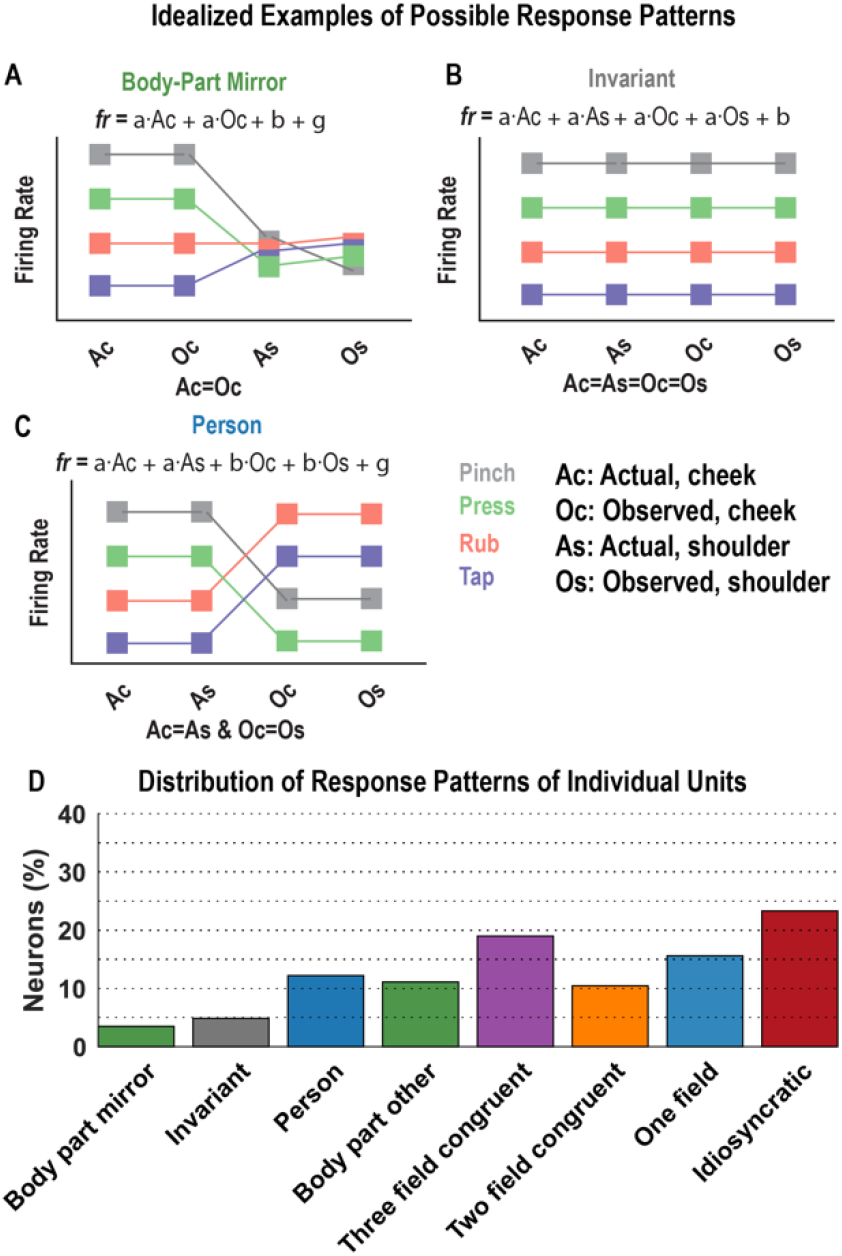
Single neurons are complex and heterogeneous. **A-C**, Schematic illustrations of three of the 51 possible linear models describing how a neuron’s firing rate response to the four touch-types (selectivity pattern, SP) is congruent or incongruent across the four sensory fields (Ac, As, Oc, Os, see legend). A more complete description can be found in Fig. S3. **D**, Histogram showing the percentage of PPC neurons that behaved according to each category of models (see also Figure S3–5). Ac, actual cheek touch; As, actual shoulder touch; Oc, observed cheek touch, Os, observed shoulder touch; fr. firing rate

The PPC population was heterogeneous, composed of many complex patterns of congruency across sensory fields. With the inclusion of the additional touch-types, only 3% of neurons show specificity and congruency for coding the body location that was touched (compared to 12% in the simple task, Fig. 1D). This result highlights the fragility of single unit mirror responses as we expand the paradigm to include a broader diversity of stimuli.

#### Population-level neural subspaces mediate the generalizability of tactile information

The mirror mechanism is proposed to link what we see with what we intend or feel. Based on our previous data within this PPC substrate, we hypothesized that the mirror mechanism is one manifestation of a broader computational strategy by which PPC neurons generalize across task dimensions (behavioral contexts). To test this hypothesis, we train a linear model to identify a neural subspace that discriminates values along one dimension for fixed values of the other two dimensions. Then, we test whether this subspace allows similar discrimination for alternate values of the fixed dimensions. For example, we train the model to discriminate touch-types (dimension 1, pinch versus press), for fixed body part (dimension 2, e.g., cheek) and person (dimension 3, e.g., NS) and test the ability to discriminate touch-type when switching body part, person, or both body part and person (Figure 4A). This analysis provides a population-level generalization of the basic mirror neuron test, testing for both specificity (operationalized by finding the population response that discriminates between two conditions) and congruency (testing whether this population response is congruent between (generalizes across) self and other, or potentially other task dimensions.) If the mirror-mechanism is the dominant motif that determines population-level encoding, then we would predict preferential congruency/generalization when matching body locations and touch-types between self and other. Otherwise, under a more general mechanism for shared encoding, we would expect generalizable information to be a ubiquitous phenomenon.

**Figure 4.**
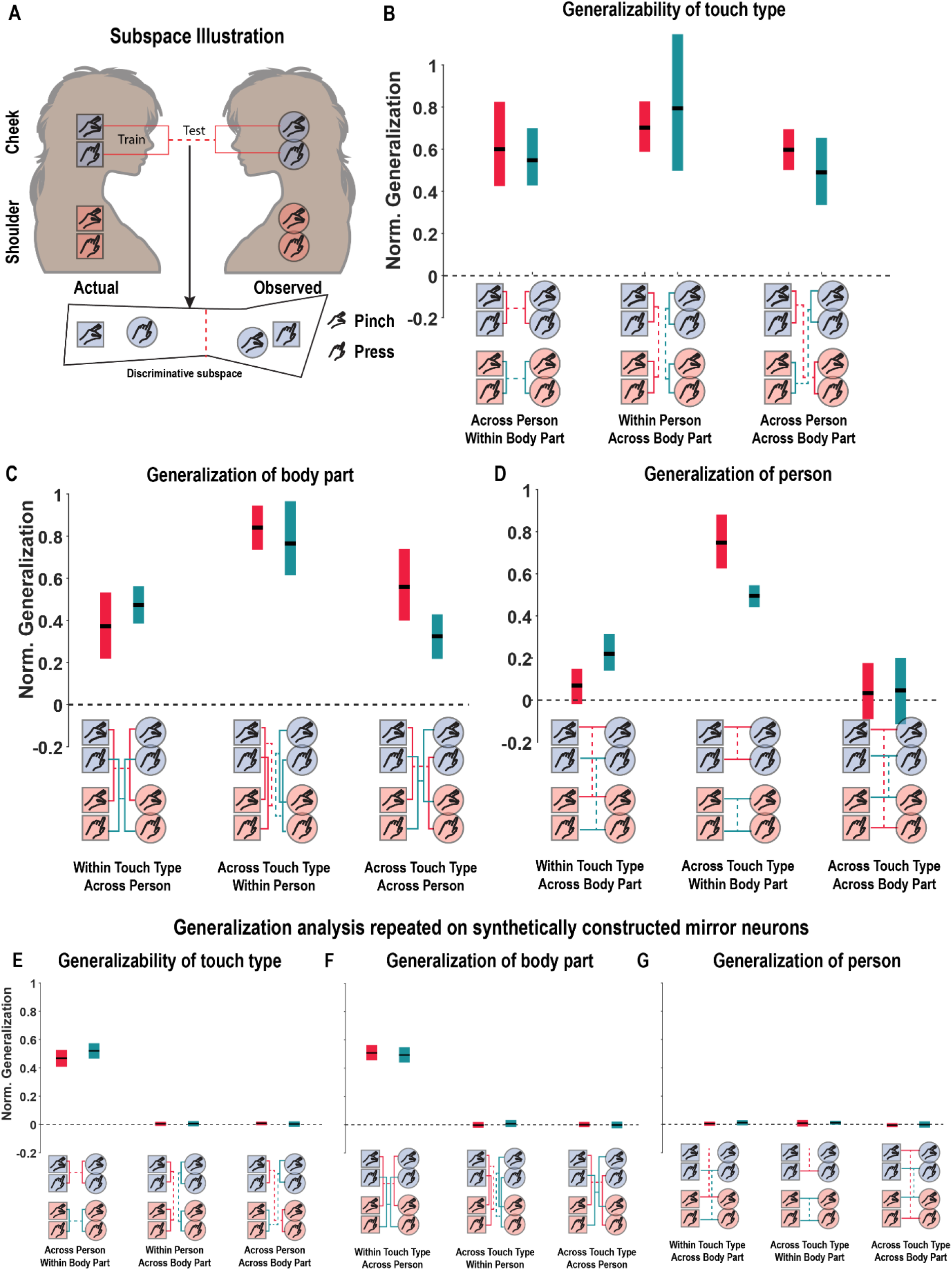
Population-level subspaces mediate the generalization of tactile information. **A**, Schematic illustration of the subspace analysis to test the generalizability of tactile information. In this example, we learn a linear mapping that discriminates actual pinch and press to the participant’s cheek from population neural activity. We then test how well the mapping is able to discriminate data collected while the participant observes pinch and press to the experimenter’s cheek. Generalization is quantified by measuring the Mahalanobis distance between conditions in the observed (test) data normalized by the distance between conditions in the actual touch (training) data. **B**, Results of the subspace analysis when testing how touch-type information generalizes across the other two dimensions: body part and person. The normalized generalization (y-axis) is shown for each tested subspace (x-axis). On the x-axis below each group of bars is a condensed schematic (from panel A), showing the train and test pairs. The red and green lines illustrate two separate but related tests for generalizability. The bars show the mean generalization (horizontal black line) ± 95% CI computed across sessions. **C**, Similar to panel B, except here the generalizability of body part information is being tested across touch type and person. **D**, Similar to panels B and C, except here the generalizability of person information is being tested across body part and touch type. Norm, normalized. **E-F**, Similar to B-D but for synthetically generated neural data.

The results of this analysis are shown in Figures 4B-D. Generalization results are scaled such that the distance between the two training conditions is equal to one. Thus a value of one for the test data would indicate perfect generalization. Positive values less than one indicate imperfect but significant generalization. A value of zero would indicate no generalization. In Figure 4B we show generalization results when discrimination subspaces are built around two of the touch-types, pinch and press. In the left panel, we show the discriminable population response that distinguishes pinch from press generalizes from self to other, both for the cheek (red) and shoulder (green). This is as expected from the mirror hypothesis. However, the response pattern generalized even more strongly when comparing within the participant’s own body (red) or the body of the actor (green) (Figure 4B, middle) and generalized equally well when mismatching body parts across person (Figure 4B, right). The logical framework of mirror neurons would not work in many of these cases since it would suggest that we understand the sensory experiences of our shoulder by simulating them within our cheek neurons (or vice versa). Alternatively, at a population level, it could suggest that touch-type is encoded as a generalizable property that can be equally well applied across self and others or different body segments.

We found a similar pattern of results when building the discrimination subspaces around body part (Figure 4C). In the left panel, we see that the discriminable population response that distinguishes observed pinches to the cheek and shoulder generalizes from self to other (red) and similarly for presses (green). However, again, we find that generalization is equally strong across the other comparisons (Figure 4C middle and right).

Interestingly, Figure 4D shows a fundamentally different pattern, with generalization in some contexts but not all. In other words, touch-type (4B) and body location dimensions (4C) are encoded in ways that generalize across all dimensions, while the person dimension only generalizes to an appreciable degree when preserving body location (middle panel). Such preferential encoding fits with the known functions of our cortical implant site (see discussion).

To assist interpretation, we repeat the analysis with a population of synthetically generated mirror neurons (Figure 4, E-G). The synthetic mirror neurons were designed with previously reported tuning complexities (see methods: Synthetic mirror neurons). The results demonstrate generalization only for matched conditions across self and other and does not support ubiquitous generalization, unlike our PPC population.

#### Architecture of knowledge representation in human PPC: structured compositionality

Our finding of universal generalization suggests that basic aspects of touch are encoded as generalizable properties that apply across multiple contexts. We, therefore, hypothesized that experienced and observed sensations are represented through a combination of simple primitive elemental features encoded by the population. We used a demixed-principal components analysis (dPCA) to visualize and quantify how much of the population-level response could be understood as being composed of a linear combination of these elemental features (e.g., body part or touch type). The dPCA analysis decomposed the population response across all test conditions into the independent factors (body part, touch type, and person) interaction effects (body part*person, body part*touch type, and person*touch type), and a term accounting for a shared neural response to any form of touch. Large contributions from the independent factors would be consistent with compositionality: that multi-faceted touch representations can be described as combinations of their basic-level components. The interaction terms capture the portion of the response that shows a higher degree of specificity. For example, the body part * touch-type interaction captures the portion of the neural response that codes for a particular combination of these two variables invariant to or generalizable across the third variable (e.g., pinches to the cheek, whether experienced or observed).

The largest percentage of the population variance relates to the shared neural response to any form of touch (Figure 5A). This response is difficult to interpret but may relate to factors like general behavioral engagement or encoding that some form of touch has occurred. Otherwise, the three main effects account for the bulk of the variance, with larger contributions from body part and touch type as consistent with Figure 4D.

**Figure 5.**
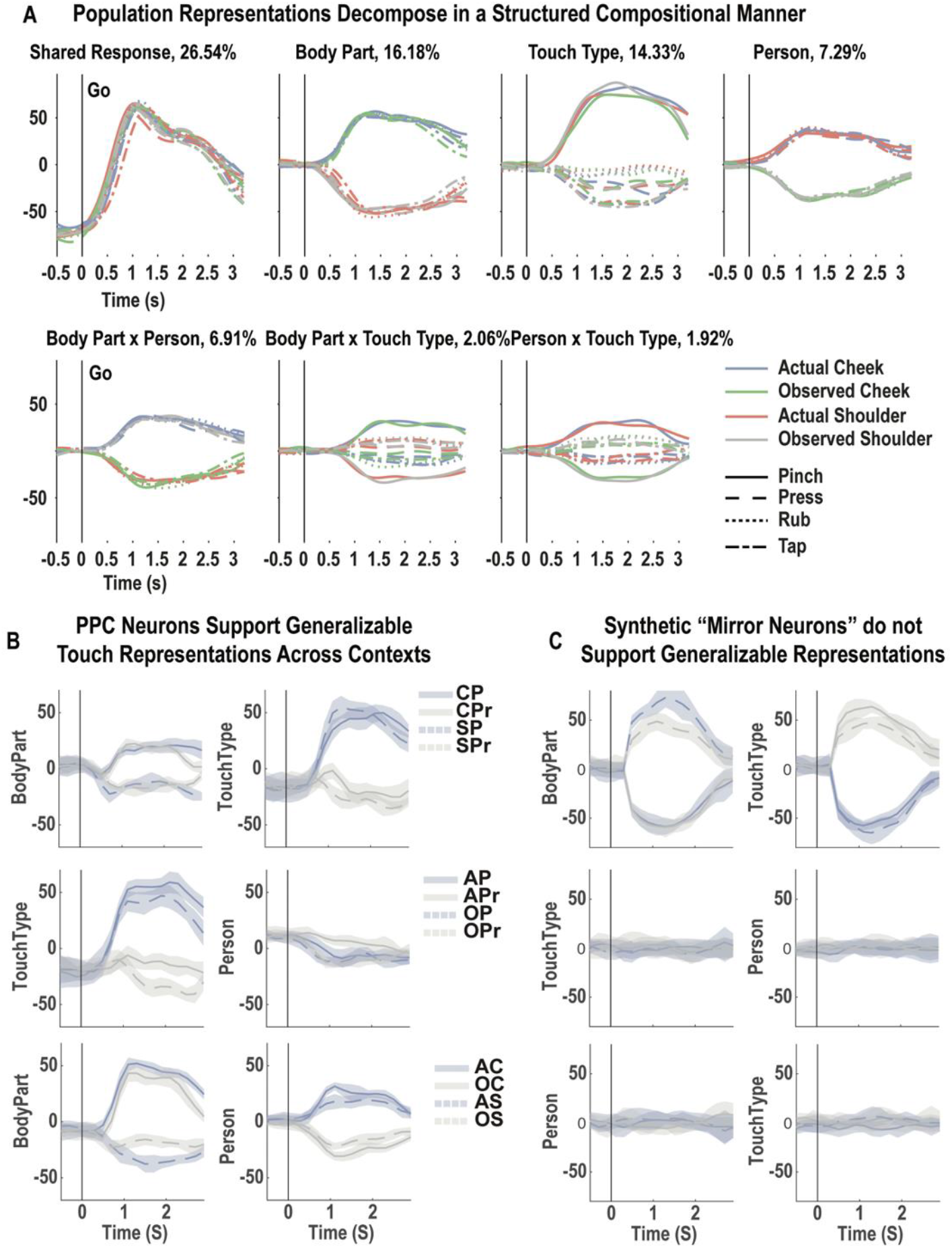
Compositional architecture of PPC responses to touch. **A**, Results of a demixed principal component analysis (dPCA) performed on the population data. Each panel shows the linear projection of the neural population response onto an axis that describes how neural firing relates to task variables, along with the percentage of the population variance that is explained by the task variable. In each panel, the projection is shown as a function of time (x-axis), along a dimensionless y-axis, coded by color and line type as labeled. **B**, Generalizability of learned neural subspaces. Each row shows the results of a dPCA in which the population representations of four conditions (in the legend) were decomposed along two dimensions (labeled y-axis; each panel) for a fixed value of the third dimension. The learned mapping was tested on the held-out value of the third dimension. The mean value across sessions (and both directions of the third dimension) ± 95% CI for each condition is shown, as a function of time (x-axis). **C**, Same analysis as in B, except performed on synthetic mirror neurons. CP, cheek pinch; CPr, cheek press; SP, shoulder pinch; SPr, shoulder press; AP, actual pinch; APr, actual press; OP, observed pinch; OPr, observed press; AC, actual cheek; OC, observed cheek; AS, actual shoulder; OS, observed shoulder; s, seconds.

To ensure that dPCA is capturing generalizable features of the neural population code, we performed a generalization test of the dPCA components. To this end, we performed a two-factor dPCA on two dimensions of the data for a single level of the third dimension. We then applied the learned mapping to the held-out level of the third dimension. The results are shown in Figure 5B for each possible combination of the two-factor dPCA. The top row tests generalization across levels of person for the two-factor body part x touch-type dPCA. The middle row tests generalization across levels of body part for the two-factor touch-type x person dPCA. The bottom row tests generalization across levels of touch type for the two-factor body part x person dPCA. As expected, we find that the main components of body part and touch type generalize, consistent with the hypothesis that PPC populations encode generalizable basic components of sensations. Similar to Figure 4, the person dimension is qualitatively different, only showing generalization across levels of touch-type, but not body-part. Finally, we repeated this dPCA analysis for our synthetic mirror neurons. As shown in Figure 5C, mirror neurons enable generalization across self and other (top row) but not for any other split of the data.

## DISCUSSION

### Mirror neurons reconsidered

Mirror neurons are the foundation of an influential theory for how we understand the actions and experiences of others (*2*). At a broad level, the mirror hypothesis claims that neurons within high-level regions responsible for planning our own motor behavior or processing bodily sensations or emotions are also involved in understanding the intentions and experiences of others. Our data are consistent with this view. The mirror hypothesis also proposes a particular mechanism by which understanding is achieved: the “mirror mechanism” (*2, 4*). In this view, individual neurons in higher-order motor cortices encode action goals, or analogous signals in other modalities (*4*). These neurons are assumed to be imbued with meaning under the assumption that we must understand our own actions or experiences. Thus, by extending access to these neurons to what we observe, the mirror mechanism can directly impart understanding (*2*).

Our results argue against this view by demonstrating: 1) that single neurons exhibiting mirror-like properties become increasingly rare as the complexity of the task increases suggesting the theory cannot scale to real-world complexity and 2) that neural population behavior is better described by shared representations that can be decomposed into basic building blocks than specifically enabling mirroring between self and other. To the first point, a number of recent studies have shown that mirroring-like behavior can be mediated by populations of neurons (*9, 22–24*), with contributions from neurons that do not directly exhibit the congruency associated with classic mirror neurons(*9*). However, the population extension of mirror neurons cannot address many of the core critiques of the theory (*6, 7*). Instead, per point two, our basic view connects more directly with developing theories of how human-like cognition relies on the nature of mental representations, outlined next.

### Potential basis for cognitive models of the world

A recent branch of cognitive neuroscience has proposed that human-like learning and thinking are primarily built on the internal models we construct of the world.(*3, 10*) These models play a ubiquitous role, providing a common substrate to inform our perception, cognition, imagination, and planning. The neural basis for this cognition through internal models framework remains largely unexplored, though preliminary neuroimaging evidence points to a role for PPC (*10, 25*). Our results, demonstrating shared representations that can be decomposed into basic building blocks, support this computational architecture, and provide preliminary insight into its neural implementation within human PPC. This framework provides a unifying account of many of our recent results, suggesting that language, imagination, planning, and perception tap into the same underlying shared internal models.(*18, 20, 26, 27*) Understanding our neural results in this framework also helps to address limitations associated with mirror neurons, as discussed below.

What does our data add to the cognition through internal models framework? First, the neural populations that underly our putative internal models are consistent with grounded or embodied notions of cognition. The tactile responses in the current study have parametrically encoded tactile receptive fields that activate within 60ms of physical contact (*28*), consistent with a bottom-up (sensory-driven) role in tactile processing. These responses can be contrasted with the highly-selective and long-latency (260-400ms) responses reported for “concept cells” within the medial temporal lobe (*29*). The fact that tactile imagery (*28*) and observation engage sensory-like populations suggests that tactile cognition is intricately tied to our somatosensory experiences and argues against purely “symbolist” views of cognition (*30*). These tactile responses are likely not raw representations of sensory inputs: neural responses in anterior regions of the PPC are consistent with state estimators that compensate for sensory delays and merge visual and somatosensory inputs (*31, 32*). Our results suggest that neural populations that help estimate the state of one’s own body may provide inductive biases that constrain and shape our cognitive understanding to be consistent with our own body knowledge.

Second, within a local neural population, the compositional nature of the population response is structured: tactile variables related to touch type and body location generalize for any split of the data, consistent with establishing a compositional basis, while the identity of who is being touched does not (Figure 4, 5). Presumably, models around identity are constructed in varied regions of the temporal cortex, including the medial temporal cortex (*33*) and the temporal-parietal junction (*34*). Overall, this pattern of results is consistent with systems-level architectures that construct understanding through the interplay of diverse but interconnected regions (*35–40*) and suggest that the state of the world is encoded as a distributed population code within and across brain regions.

### Practical differences with the mirror neuron account: generalization and temporal dependence

By the mirror account, single neurons encode our own action goals, and the mirror mechanism extends access to these representations to observation (*41*). As stated above, such a mechanism does not allow for understanding action goals outside our own repertoire. In our account, high-level cortical regions encode representational building blocks that can be combined in novel ways to understand novel stimuli. For example, we would predict that if our participant was asked to imagine what it would feel like if we pinched her tail (something clearly outside the participant’s direct experience as she lacks a tail), we would find that the neural subspace associated with pinches would be activated along with cortical representations of tails, presumably derived from regions of the temporal cortex (*42*).

The mirror account assumes a temporal dependence: individuals learn motor representations that can subsequently be accessed during observation (*2*). In our formulation, such temporal precedence is not necessary. Individuals can form representational building blocks using any possible information source. To use an example from Patricia Churchland, even if I have never milked a cow, I can readily build understanding by watching someone else milk a cow. The underlying neural representations can then help me quickly understand milking a goat or help inform my own attempts at milking a cow (*43*). No doubt, our actions and experiences provide unique and unreproducible forms of knowledge. Just as the description of a sunset cannot replace witnessing a sunset, it is likely that observing pinches cannot replace the experience of having been pinched. Thus, our own experiences can change the nature of our internal representations but are not a prerequisite to form these representations. As evidence, even individuals with congenital somatosensory deprivation can form relatively natural internal representations of the body (*44*).

#### Compositionality

Compositionality captures the basic idea that we construct representations through a combination of more primitive components. For example, a car can be encoded as a combination of wheels, body, engine, seating, steering mechanism, etc. The same elements can be recombined in different ways to form related representations, such as a bus or motorcycle. Two primary approaches have been used to test for compositionality in neural populations: matrix factorization and parts-whole based approaches. The nature of our stimuli naturally lent itself to the matrix-factorization based approach (see methods, task description) and may be necessary to ensure that adequate context. For example, the pantomimed gesture of two fingers pressing together may not equate to a “pinch.” To this point, observation of motor movements devoid of goals is insufficient to drive action observation neurons (*4*). Nonetheless, questions about what constitutes a sufficient stimulus and delving further into the mechanisms of compositionality are exciting directions for future studies.

Compositionality does not imply a specific neural architecture. For example, “concept cells,” neurons described in the medial temporal lobe (*45*) that respond to a preferred stimulus (e.g. a particular individual) independent of sensory modality or presentation details (e.g., image, written word, sound) can form a compositional basis. For example, the concept of “Star Wars” may be formed by an ensemble of cells encoding subconcepts such as “Luke Skywalker,” “Darth Vader,” etc (*46*). We have previously described a partially-mixed architecture. Unlike concept cells, PPC neurons respond to many diverse stimuli in seemingly random ways at an individual cell level. For example, a cell encoding hand movements was as likely as not to encode a shoulder movement (*20, 47*). Nonetheless, neurons exhibit clear structure at a population level, forming associations between related variables, consistent with this work. One possibility is that compositionality built on partially-mixed representations helps embody or tie our understanding to our lived experiences.

### Relationship to alternative accounts of mirror neurons

Following critiques of the mirror hypothesis (*7, 48, 49*), alternative explanations for cells that fire to both performed and observed movements have focused on a role for the visual guidance of movement, e.g., by mediating motor imitation, observational learning, or planning in response to the actions of others (*41*). These are compelling as animals clearly use such observation to guide their motor behavior. It is less clear how well such explanations can account for our data: There is no simple corollary for generating an endogenous sensory experience in response to the sensory experiences of others. In our view, the compositional building blocks provide useful representations that can inform all relevant aspects of behavior. In the motor domain, this can include e.g. action understanding, but also guiding motor behavior based on the actions of others. Interestingly, the degree of population-level similarity between executed and observed actions appears smaller than what we have found in the sensory domain (*41*). One intriguing hypothesis is that the number of possible ways observed actions can inform our cognition and behavior (e.g., understanding, imitation, learning, motor planning) is substantially larger than the sensory case leading to more multi-faceted neural responses that limit gross measures of population similarity.

### Relevance to BMI

Numerous clinical trials have shown that individuals with paralysis can use signals from motor regions of the brain to control external devices, such as robotic limbs or computer cursors (*18, 50*). The underlying brain signals are low-dimensional and roughly encode movement direction smoothly, enabling researchers to collect sufficient data to train a decoding algorithm in a few minutes. Future BMIs could decode high-level concepts, visual imagery, or emotional state. The dimensionality of these signals (e.g., the space of all mental images) is far larger than basic movements. However, if these high-dimensional datasets are encoded using generalizable relatively low-dimensional basis sets, then the ability to read out these high-dimensional signals may be tractable. To this end, proof-of-concept studies have already demonstrated the ability to decode high-fidelity faces or the semantic content of visual scenes from rich low-dimensional basis sets (*17, 51, 52*). The current study suggests that complex high-level concepts can be constructed as the linear superposition of basic level representations (e.g., touch type or body location) that are decodable from limited datasets. This offers hope that future BCIs can communicate the contents of our minds more directly, providing truly novel forms of communication.

## Acknowledgments

We thank NS for her bravery and hard work in making this work possible.

## Funding

National Institute of Health grant R01EY015545 (RAA, TA, NP)

Conte Center for Social Decision Making at Caltech grant P50MH094258 (RAA)

Tianqiao and Chrissy Chen Brain-machine Interface Center at Caltech (RAA, TA)

Boswell Foundation (RAA)

## Author contributions

Conceptualization: TA

Methodology: TA, SC

Investigation: TA, CZ

Formal Analysis: TA, SC

Visualization: TA, SC

Funding acquisition: TA, NP, RAA

Project administration: TA, RAA

Supervision: TA, NP, RAA

Writing – original draft: TA, SC

Writing – review & editing: TA, SC, RAA

## Data and materials availability

The datasets analyzed for this manuscript will be shared upon reasonable request.

## Materials and Methods

### 1. Subject details

All data were recorded from NS, a 62-year-old tetraplegic female participating in a brain-machine interface (BMI) clinical trial. She has a high-cervical spinal cord injury (SCI) between cervical levels three and four, sustained approximately 10 years prior to the study, and with no preserved sensory or motor function below the shoulder. She was implanted with two 96-channel Neuroport Arrays (Blackrock microsystems model numbers 4382 and 4383) 6 years post-injury, in the left hemisphere. Informed consent was obtained, and she understood the nature, objectives, and potential risks of the surgical procedure and the subsequent clinical studies. All procedures were approved by the Institutional Review Boards (IRBs) at the California Institute of Technology (IRB #18-0401), the University of California, Los Angeles (IRB #13-000576-AM-00027), and Casa Colina Hospital and Centers for Healthcare (IRB #00002372).

### 2. Experimental setup

All experiments were conducted at Casa Colina Hospital and Centers for Healthcare. NS was seated in a motorized wheelchair in a well-lit room. A 27-inch LCD monitor was positioned behind NS (visible to the experimenters but not to NS) to cue the experimenters when to deliver tactile stimuli. Cue presentation was controlled by the psychophysics toolbox (Brainard, 1997) for MATLAB (MathWorks).^83^

### 3 Physiological recordings

NS was implanted with one Neuroport array at the junction of the intraparietal sulcus (IPS) and postcentral sulcus (PCS), a region we refer to as PC-IP.^34^ The other array was implanted in the left superior parietal lobule (SPL). Following surgery, the SPL implant did not function. Only data recorded from PC-IP were used in this study. Both arrays were explanted approximately two years after data in this study were collected.

Neural activity recorded from the array was amplified, digitized, and sampled at 30 kHz using a neural signal processor. This system has received Food and Drug Administration (FDA) clearance for <30 days of recordings. We received an investigational device exemption (IDE) from the FDA (IDE #G120096, G120287) to extend the implant duration for the purposes of the BMI clinical study.

Putative neuron action potentials were detected at threshold crossings of −3.5 times the root-mean-square of the high-pass filtered (250 Hz full bandwidth signal. Each waveform was made of 48 samples (1.6 ms), with 10 samples prior to triggering and 38 samples after. Single- and multi-unit activity was sorted using Gaussian mixture modeling on the first three principal components of the detected waveforms^35^. To minimize noise-related effects, we used, as selection criteria, a mean firing rate greater than 0.5 Hz and signal to noise ratio (SNR) >0.5.

### 4 Task procedures

#### 1 Basic sensory mirroring task (BSMT; relevant for Figure 1)

This task was performed to establish the shared responsiveness of PPC neurons to ***felt*** and ***observed*** touch. NS sat facing an experimenter (actor). One experimenter stood behind the actor, and another behind NS. The task involved touch to one of two body parts (cheek, shoulder), to one of two persons (subject, or actor). Touch was provided as rubs performed bilaterally by the experimenter standing behind the person being stimulated, at approximately 2 rubs per second, for 3 seconds. Cheek touches were rubs parallel to the jawline (from cheekbone to chin and back again). Shoulder touches were rubs along the top of the shoulder, from near the neck to the outside of the shoulder and back. The task was performed on 6 individual recording sessions, with 10 trials per condition. In all, 805 units were recorded, of which 756 met the selection criteria.

#### 2 Multidimensional sensory mirroring task (MSMT; relevant for all Figures except 1)

This task was performed to understand mechanisms by which neural information is shared across populations of PPC neurons to support ***felt*** and ***observed*** touch. The basic setup was like the previous task. Here, however, we manipulated three dimensions: 2 body parts (cheek, shoulder), provided to 2 persons (NS, actor), in one of four touch-types (pinch, press, rub, tap). As in the BSMT, touch stimuli were provided at approximately 2 per second, for 3 seconds. Rubs were as described. Pinches were performed in a non-painful manner with the thumb, index, and middle fingers. Presses were performed with the index and middle fingers and taps by the tips of the index and middle fingers. Prior to performing the experimental session, we verified that the participant was able to differentiate the different stimuli, whether observed or felt. This task was performed on 8 recording sessions, with 10 trials per condition. In all 806 units were recorded, of which 741 met the selection criteria.

The use of three factors was essential to enable testing for compositionality in the neural code. There are two primary approaches that have been used to test for compositionality in neural populations: matrix factorization and parts-whole based approaches. In the matrix factorization approach, population activity is measured while the participant views complex stimuli composed of varied combinations of different constituent elements (*13, 14, 16, 17*). The resulting activity patterns are then subjected to some form of matrix decomposition to test whether the stimuli are explainable as combinations of the constituent elements. For example, a stimulus set may consist of images of young men and women and old men and women. The resulting brain activity would then be decomposed to test whether neural responses can be represented as combinations of age and gender dimensions. In the parts-whole approach, population activity is measured while the participant views stimuli of the parts and the whole separately (*15*). Brain responses evoked by the parts are then combined to see if they predict the brain response to the whole. For example, a stimulus set may consist of the words “old,” “man,” and “grandpa.” Response to “old” and “man” are then summed to see if they predict “grandpa.” This latter approach has intuitive appeal as it has a direct relationship to the underlying theory – the whole is the sum of its parts. However, representing the parts is not trivial for certain classes of stimuli. One cannot simply show an image of “old”; there must be an embodiment – an image of something old. Additionally, it is unclear whether parts, devoid of their surrounding context, would or should be processed as such. For example, the pantomimed gesture of two fingers pressing together may not equate to a “pinch.” To this point, observation of motor movements devoid of goals are insufficient to drive action observation neurons (*4*). These latter cases naturally lend themselves to the matrix factorization approach – e.g. to show stimuli of the “old something” and use an appropriate analysis technique to separate the neural signature of “old” from the neural signature of the “something.” This approach carries the possibility of overfitting the data, finding a separation that is not a true generalizable feature of the data. To mitigate against this possibility, it is essential to record a dataset that allows any identified structure to be validated against data that was not used to train the algorithm. Such an approach requires a sufficient number of factors, at minimum three, so that the matrix decomposition can across at minimum two factors and tested across varied levels of the third factor.

### Quantification and statistical analysis

#### 5.1 Linear analysis (relevant for Figure 1B, Figure 2A)

For each unit, we fit a linear model describing its firing rate as a function of response to each test condition. Response was defined as the mean firing rate between 0.5 after onset of the stimulus phase and ending 0.5 s thereafter. These times were chosen to correspond to the period of time during active tactile stimulation, offset to account for experimenter delays in presenting the stimulus. The baseline was defined as the neural firing rate during the 1 s prior to stimulus presentations. The linear model was computed as:

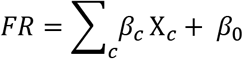

Where *FR* is the firing rate, X_*c*_ is the vector of indicator variables for test condition *c*, *β_c_* is the estimated scalar weighting coefficient for each condition, and *β*_0_ is a constant offset term. A neuron was considered responsive to a particular condition if the t-statistic for its associated beta coefficient was significant (*p*<0.05, false discovery rate (FDR) corrected for multiple comparisons).

#### 5.2 Discriminability index (relevant for Figure 1C, Figure 2B)

To quantify how well neural activity can be discriminated from baseline (pre-stimulus) activity, we used a cross-validated mahalonbis distance measure. As with the linear analysis described above, the stimulation phase window was defined as 0.5 after onset of the stimulus phase and ending 0.5 s thereafter, and baseline was defined as the 1 s prior to stimulus presentation). The firing rate of all recorded neurons was concatenated into a vector, denoted by A. The firing rate of each neuron during the baseline phase was similarly concatenated to form a vector, denoted by *B*. Next, a non-dimensional distance was computed as:

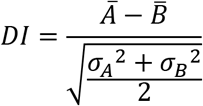

Where 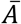 is the mean of the firing rate vector 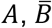 is the mean of the firing rate vector *B*, *σ_A_* is the standard deviation of the vector *A*, and *σ_B_* is the standard deviation of the vector *B*.

#### 5.3 Time-resolved classification (relevant for Figure 2C)

Classification was performed using linear discriminant analysis (LDA) with the following parameter choices: (1) only the mean firing rates differ for unit activity in response to each test condition (covariance of the normal distributions are the same for each condition) and (2) firing rates for each unit are independent (covariance of the normal distribution is diagonal). The classifier took as input a matrix of firing rates for all sorted units. The analysis was not limited to significantly modulated units to avoid ‘peeking’ effects.^84^ The analysis was performed independently for each recording session, and results were then averaged across days. In **Figure 2C**, this analysis was performed in a sliding-time window manner (300 ms each window, stepped at 10 ms intervals), beginning 0.5 s prior to the stimulation onset. Classification performance is reported as the prediction accuracy of a stratified leave-one-out cross-validation analysis.

#### 5.4 Correlation (relevant for Figure 1F)

We performed cross-validated correlation to compare the neural representations of various test conditions (stimulus presentations) against each other in a pairwise manner. We quantified the neural representations as a vector of firing rates, one vector for each condition with each vector element summarizing the response of an individual unit. Neural activity was summarized as the mean firing rate during the stimulation phase window, defined as before (0.5 s after onset of the stimulus phase to 0.5 s after it ended). Firing rate vectors were constructed by averaging the responses across 50–50 splits of trial repetitions. The mean responses across different splits were correlated within and across conditions, then the splits were regenerated, and the correlation computed 250 times. The within-condition correlations assist in our interpretation of the across-sensory field correlations by allowing us to quantify the theoretical maxima of the similarity measure (e.g., if the within-condition correlation is measured at 0.6, then an across condition of 0.6 suggests the maximal level of similarity as allowed by the trial-to-trial variability of the signal).

#### 5.5. Event-related averages (relevant for Figure 1E, Figure 2D, Figure 2E, Figure 2F, Figure S2)

For each unit, neural activity was averaged within 750 ms intervals starting from 0.5 s prior stimulation onset, stepping to 2.5 s after, in 100 ms step intervals. Responses were grouped by condition, and a mean and standard error on the mean (SEM) were computed for each time window and for each condition.

#### 5.6 Modeling the single neuron response properties to various test conditions (BSMT) (relevant for Figure 1D)

This analysis was performed to understand how individual neurons responded to four formats: actual cheek touch (Ac), actual shoulder touch (As), observed cheek touch (Oc), and observed shoulder touch (Os). Various possibilities exist. For example, the neuron might respond to actual touch to both body parts but not to any observed touch. Alternatively, it could respond to both actual and observed touch to the one body part but not to the other. We can model the firing rate for a given unit as:

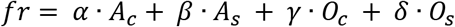

Where *fr* is the firing rate for the unit, *A_c_*, *A_s_*, *O_c_*, *O_s_* are the four formats, and *α*, *β*, *γ*, and *δ* are the weighting coefficients for each sensory field, respectively. If the unit does not respond to a sensory field, then the dot product of the unit’s weighting coefficient and the sensory field collapses to a scalar value. If a unit responds to two formats in a congruent manner, then the weighting coefficient for these two formats will be identical. For the analysis, we allowed a weighting coefficient to be either 0 or 1, such that that across 4 formats, there are a total of 16 possible models for each neuron. We fit the parameters of each of the 16 models using standard linear regression techniques (see above), and the results were compared. As selection criteria to evaluate the “best” model from all candidate models, we used the Bayesian information criterion (BIC) and cross-validated coefficient of determination (cvR^2^). The models were grouped according to four categories: **invariant** (in which the weighting coefficient was identical across all formats), **body part** specific (in which the weighting coefficient was invariant for matched body parts, but not for mismatched body parts), **person** specific (in which the weighting coefficient was invariant for touch to the same person) or **idiosyncratic** (all other combinations).

#### 5.7 Modeling the single neuron response properties to various test conditions (MSMT) (relevant for Figure 3, Figure S3, Figure S4)

This analysis is like the earlier modeling analysis for the BSMT, except it has been expanded to accommodate for more test conditions. To understand the breakdown of individual units that create the population response, we first defined four formats: ***actual*** cheek touch (Ac), ***actual*** shoulder touch (As), ***observed*** cheek touch (Oc), and ***observed*** shoulder touch (Os). An individual neuron could respond to one or more formats. If it responds to more than one sensory field, it could respond with a congruent selectivity pattern (SP; the precise pattern of responses) to each of the four touch types (pinch, press, rub, tap) within the sensory field, or with an incongruent SP. Across the four formats, the firing rate for a given unit can be described mathematically as:

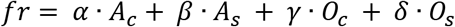

Where *fr* is the firing rate for the unit, *A_c_*, *A_s_*, *O_c_*, *O_s_* are the four formats, and *α*, *β*, *γ*, and *δ* are the weighting coefficients for each sensory field, respectively. If the unit does not respond to a sensory field, then the dot product of the unit’s weighting coefficient and the sensory field collapses to a scalar value. Within this type of a linear model, if a unit responds to formats with an identical SP, then the weighting coefficient for all those formats will have an identical weighting coefficient. In all, there are 51 unique models for all the ways in which SPs can be expressed across formats.

To determine how SPs compared across formats, we fit the parameters of each of the 51 models using standard linear regression techniques (see above), and the results were compared. As selection criteria to evaluate the “best” model from all candidate models, we used the Bayesian information criterion (BIC) and cross-validated coefficient of determination (cvR^2^). Results are summarized as the number of units that are best described by a particular model.

#### 5.8 Generalizability analysis (relevant for Figure 4)

This analysis was performed to understand how neural information belonging to one domain (e.g., body part) generalizes across another domain (e.g., touch type or person). For this analysis, we restricted the tested touch types to pinch and press only. Additional combinations of touch type, e.g. pinch and rub were tested and resulted in the same pattern of results. Restricting to two touch-types resulted in three data matched dimensions: touch type (pinch, press), body part (cheek, shoulder), and person (actual touch, or observed touch). In all, there are 8 test conditions.

We quantified the neural response to each condition as a vector of firing rates, one vector for each condition, with each element in each vector summarizing the response of an individual unit. All sorted units were used in this analysis. As in other analyses, neural activity was summarized as the mean firing rate beginning 0.5 s after onset of stimulation phase and ending 0.5 s after its conclusion. Next, we identified population-level neural subspaces that optimally differentiate between pairs of conditions (training conditions) and tested how well these subspaces differentiate between pairs of test conditions. For example, in Figure 4A, we identified a subspace that optimally differentiates between the two touch types, pinch and press, during actual touch the participant’s cheek and asked: how well does this subspace also differentiate between touch-types when observing them applied to the cheek? To create the subspaces, we linearly regressed (using partial least squares regression) the vectors for the pair of training conditions, such that the cross-validated Mahalonbis distance between the two conditions was maximized. We then used this model to project the held-out test data into the same subspace and computed the Mahalanobis distance between conditions. This computed distance was normalized to the cross-validated distance of the training data and thus the resulting metric expresses how well the test conditions are separable relative to the training conditions. In this way, the analysis is able to tell us how well a neural subspace that optimially distinguishes the training conditions is able to apply o the test conditions. A value of 1 indicates that the neural subspace that maximally separates the training data (e.g., measured during actual touch) perfectly generalizes to the held-out test data (e.g. measured during observation). This basic computation was performed in reverse as well, such that in this example, a subspace was created that optimally differentiated between ***observed*** cheek pinch and presses, and tested to identify separability between ***actual*** cheek pinches and presses. The results in both directions were averaged and recorded as the normalized generalizability of information for presentation purposes (in this example case, the generalizability of information separating touch-types across ***actual*** and ***observed*** touch). The generalizability was computed for each day independently and averaged across recording sessions. Confidence intervals were estimated using a bootstrap procedure.

#### 5.9 Generalizability analysis applied to synthetic mirror neurons (relevant for Figure 4E-G)

The generalizability analysis was performed on both real neural data, as well as a population of synthetically generated mirror neurons. We generated neurons with response properties that allowed single neurons to unambiguously associate observed and actual touches. The neurons were generated by assigning identical firing rates to actual and observed touch to a randomly chosen body part, and a randomly chosen touch type (between pinch and press). A vector of firing rates (of length 10, for ten trials) was constructed, with a mean of 1, and a standard deviation of 1.2, for that pair of conditions, and 0 for all other pairs of conditions. To reflect the inherent complexity of real-world populations recorded in mirror neuron experiments^85^, we also created neurons selective to actual touches only, observed touches only, and observed and actual touches with different response profiles. This population of synthetic neurons was then used as the dataset for the generalizability analysis (Figure 4E-G). We also ran the analyses with only the population of cells that showed congruency between observed and actual responses. Results were identical, save that measured slightly higher generalization values for the mirror conditions, as expected.

#### 5.10 Demixed principal components analysis (relevant for Figure 5)

Demixed-principal component analyses (dPCA) is an analysis technique that decomposes population neural data along user-defined neural dimensions (marginalizations) that capture variance related to task variables ^86^. This decomposition provides insight into the structure in neural data as it relates to the experimentally manipulated variables. We used all sorted units in this analysis. dPCA takes as input a matrix that describes firing rates to each of the test conditions for each trial as a function of time (i.e., all combinations of experimental variables, 16 conditions in the MSMT task). Neural activity was averaged within 750 ms intervals starting from 0.5 s prior to the onset of the stimulation phase, stepping to 0.25 s after the stimulus offset, in 50 ms step intervals, to the time of stimulus offset. In our current study, we were interested in understanding how much of the population variance was explained by independent dimensions (i.e., body-part, touch type, and person being touched) as well as by interaction terms (touch type x body-part, body-part x person, touch type x person, and touch type x person x body-part). Thus, we used all 7 possible marginalizations within this analysis.

#### 5.11 Generalizability analysis for demixed principal components analysis (relevant for Figure 5B-C)

Our interpretation of the dPCA analysis assumes that neural subspaces associated with task variables are generalizable, capturing aspects of the data that would apply to different contexts and stimuli. For example, the neural subspace associated with the main effect of body location should code for that body location whether touch is actual or observed, and for all forms of touch. In order to validate whether the discovered subspaces are indeed generalizable, we repeated the dPCA analysis, but now explicitly testing for generalizability of the different components. To this end, we trained a two-factor dPCA on multiple levels of two dimensions of the data for a single fixed value of the third dimension. We then test the subspaces using entirely held-out data, from the alternate value of the third dimension. For example, training the dPCA using data acquired during actual touch (e.g. for multiple touch types and body-locations) and applying the solution on data acquired during observed touch. If the dPCA accurately accounts for the neural variability in the held-out data, then we can interpret the subspaces as being a generalizable feature of the neural population. We performed this generalization analysis for all ways of partitioning the three dimensions into two training and one test dimension. Further, we performed the same set of tests using a synthetic mirror neuron population. The mirror neuron population was constructed identically as described above in section 5.9 expect that we gave neural response temporal dynamics. Temporal dynamics were created by assuming no neural modulation up until .5 seconds and then assuming a stationary response through the “stimulus period”.

## Supplementary Figures

**Figure S1:**
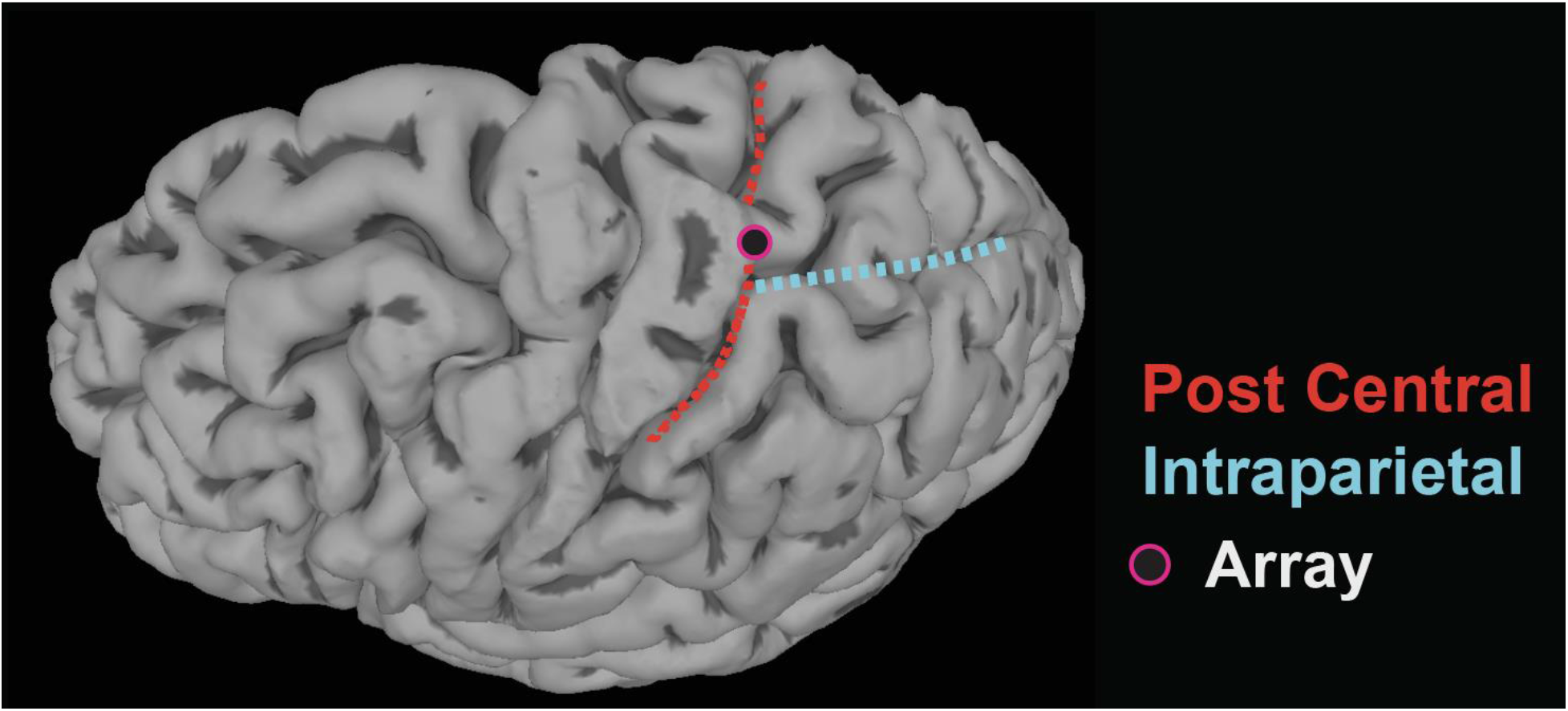
Microelectrode array implantation location. Related to Figures 1&2. Individual participant anatomy with the location of the microelectrode array implant.

**Figure S2.**
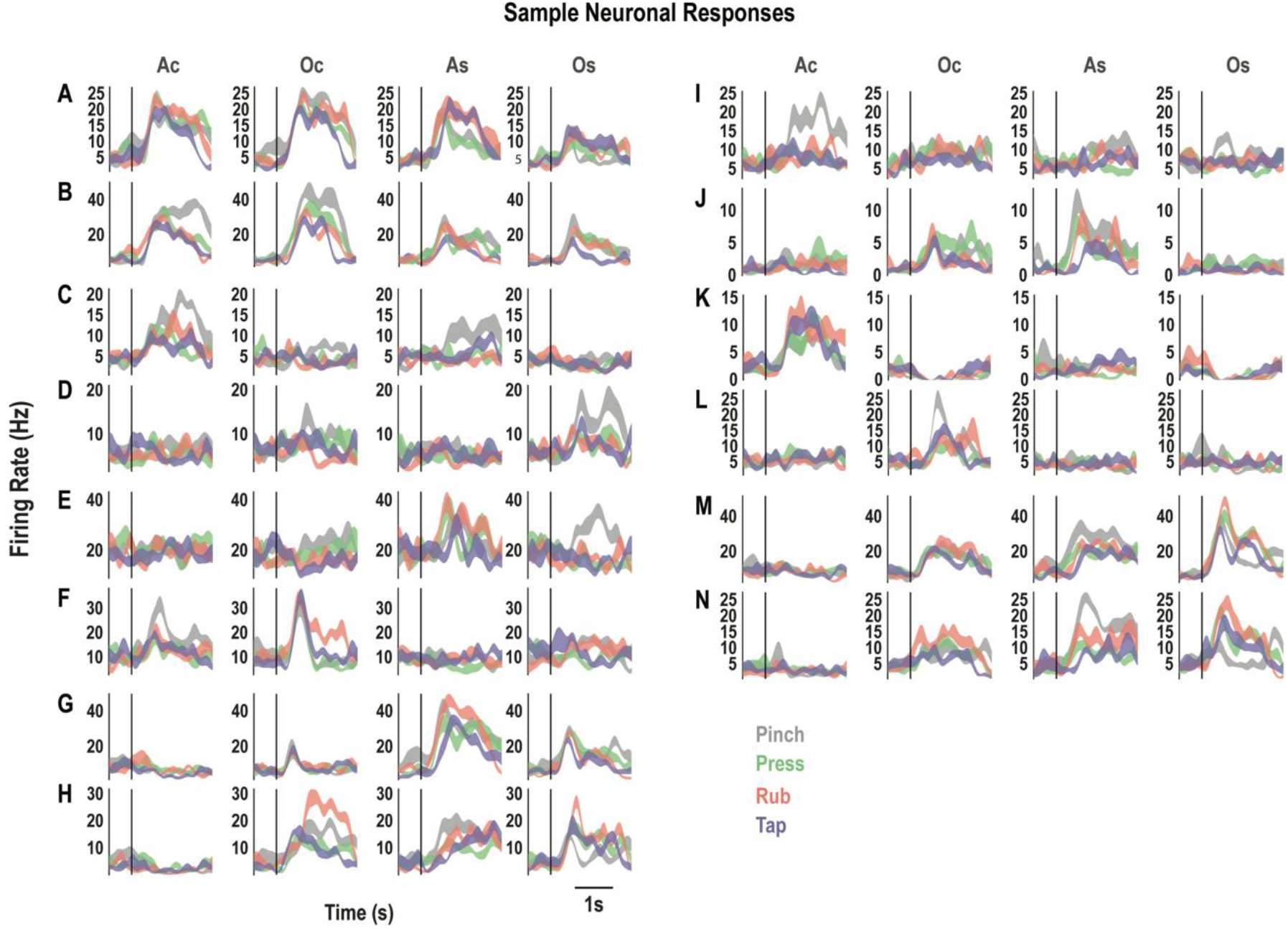
Individual neurons exhibit complex and variable response patterns. Related to figure 2. **A-N**, Firing rate as a function of time for example neurons illustrating diverse responses to tactile stimuli across the different sensory fields (columns; Ac, actual cheek touch; Oc, observed cheek touch; As, actual shoulder touch; Os, observed shoulder touch). Within each panel, the neural response to each of the four touch types is shown (colors as in legend), as the mean firing rate (y-axis) ± SEM, n=10 trials, as a function of time (x-axis). Ac, actual cheek; Oc, observed cheek; As, actual shoulder; Os, observed shoulder; Hz, hertz; s, seconds.

**Figure S3.**
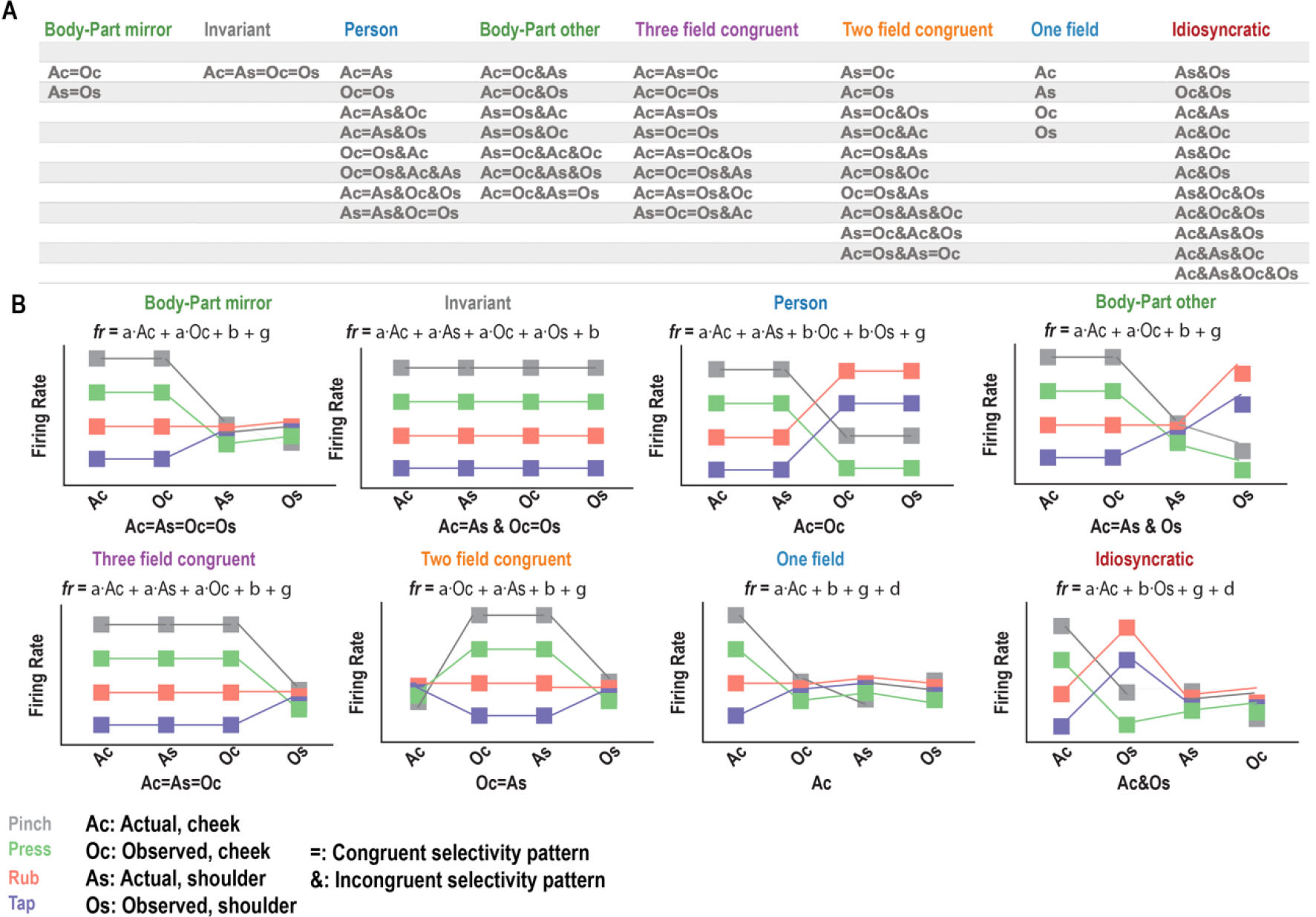
Schematic illustration describing congruency in selectivity patterns across sensory fields. Related to figure 3. **A**, All 51 possible linear models that could describe an individual neuron’s response (selectivity pattern, SP) to the four different touch types. The ‘=’ symbol denotes congruency in SP between the sensory field listed before and after the symbol, and the ‘&’ symbol denotes incongruent SPs. The 51 possible models are grouped into 8 categories for interpretative purposes, labeled accordingly. Ac, actual cheek touch; As, actual shoulder touch; Oc, observed cheek touch; Os, observed shoulder touch. **B**, Schematic illustrations of each category of the 51 possible linear models. Each panel shows one example case for the columns shown in panel A, as labeled underneath each panel. The examples illustrate cases in which the SP is congruent between actual and observed touch to the cheek but unresponsive to shoulder touch (body-part mirroring), the SP is congruent across all sensory fields (invariant), and congruent within-person (i.e., for actual cheek and actual shoulder touch) but incongruent from actual to observed touch (person). The mathematical description of each model is shown above each illustration. In the analysis, congruency is operationalized by using the same linear-model coefficients to describe responses to multiple sensory fields. Incongruency uses distinct linear model coefficients. An abbreviated name for the model is shown below each panel. The ‘=’ symbol denotes congruency in SP between the sensory field listed before and after the symbol, and the ‘&’ symbol denotes incongruent SPs across sensory fields. See figure S3 for a full list of models and examples for each category of behavior. Ac, actual cheek touch; As, actual shoulder touch; Oc, observed cheek touch, Os, observed shoulder touch; fr, firing rate. **Description of the eight summary models**: The eight models can be understood as follows: Body Part Mirror describes models in which there is unambiguous congruency between actual and observed touch, specific to a body part. Invariant describes the case in which neural responses are congruent across all sensory fields. Person describes neurons where there is a congruent response to the different body parts within an individual, but incongruency between actual and observed touch. Body part other describes neurons with congruency between actual and observed touch to one body part, but some form of incongruency between body parts thus creating ambiguity about the stimulus at the single neuron level. Three and two field congruent describes neurons showing congruency across 3 or 2 of the 4 sensory fields, but are not well-described by the other models (e.g., an example would be congruency between actual shoulder and observed cheek responses.) One field describes neurons that show discriminable responses between touch types for only one field. Idiosyncratic describes neurons that show incongruent responses to two or more sensory fields.

**Figure S4.**
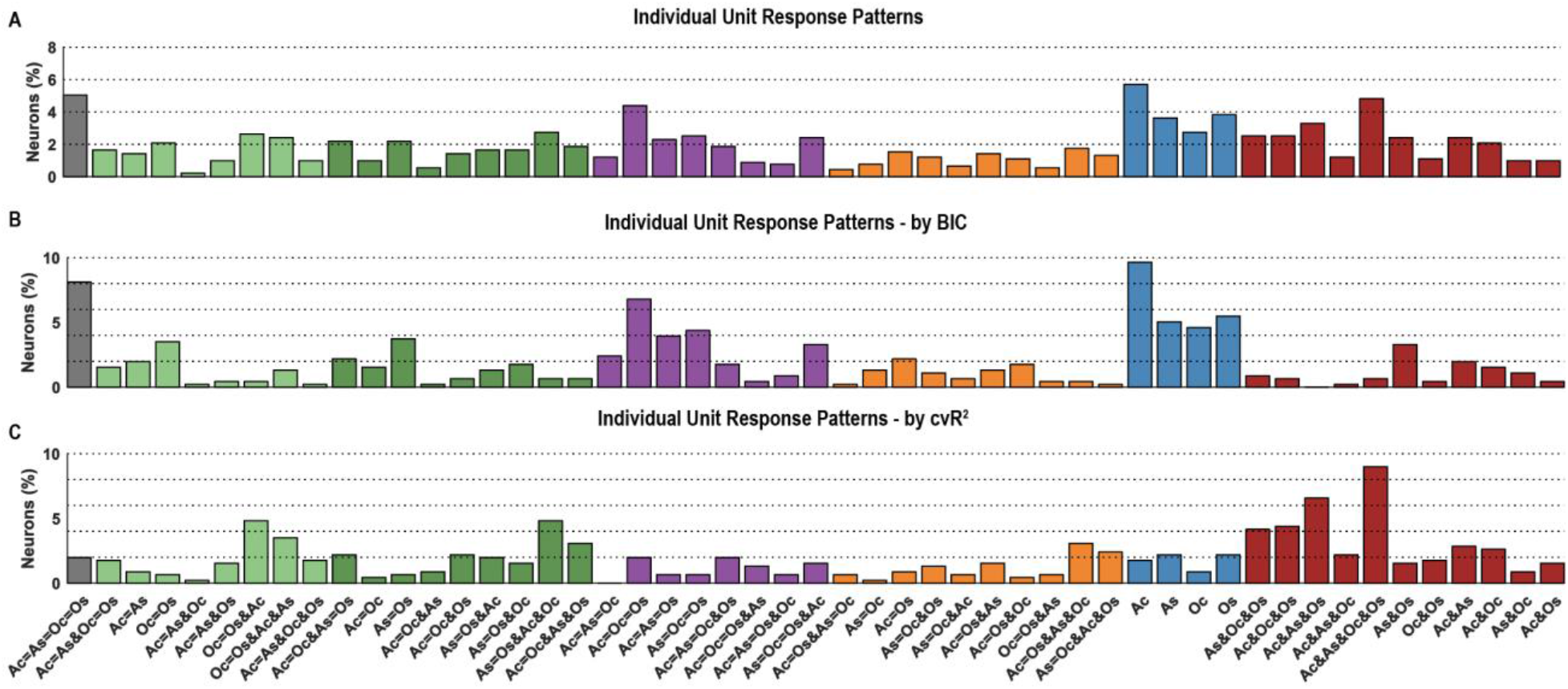
Categorization of all single-units into 51 possible patterns of correspondence across sensory fields. Related to figure 3. **A**, Histogram showing the percentage of PPC neurons that behaved according to each of the 51 possible linear models. The abbreviated form of the model is listed below each bar. As in Figure S3, the ‘=’ symbol denotes congruency in SP between the sensory fields listed before and after the symbol, and the ‘&’ symbol denotes incongruency. Bars are color-coded by category (see figure 3). Ac, actual cheek touch; As, actual shoulder touch; Oc, observed cheek touch, Os, observed shoulder touch. **B-C**, Similar to A except categorizations are shown for Bayesian information criteria (BIC) and the cross-validated coefficient of determination (cvR^2^) separately.

**Figure S5.**
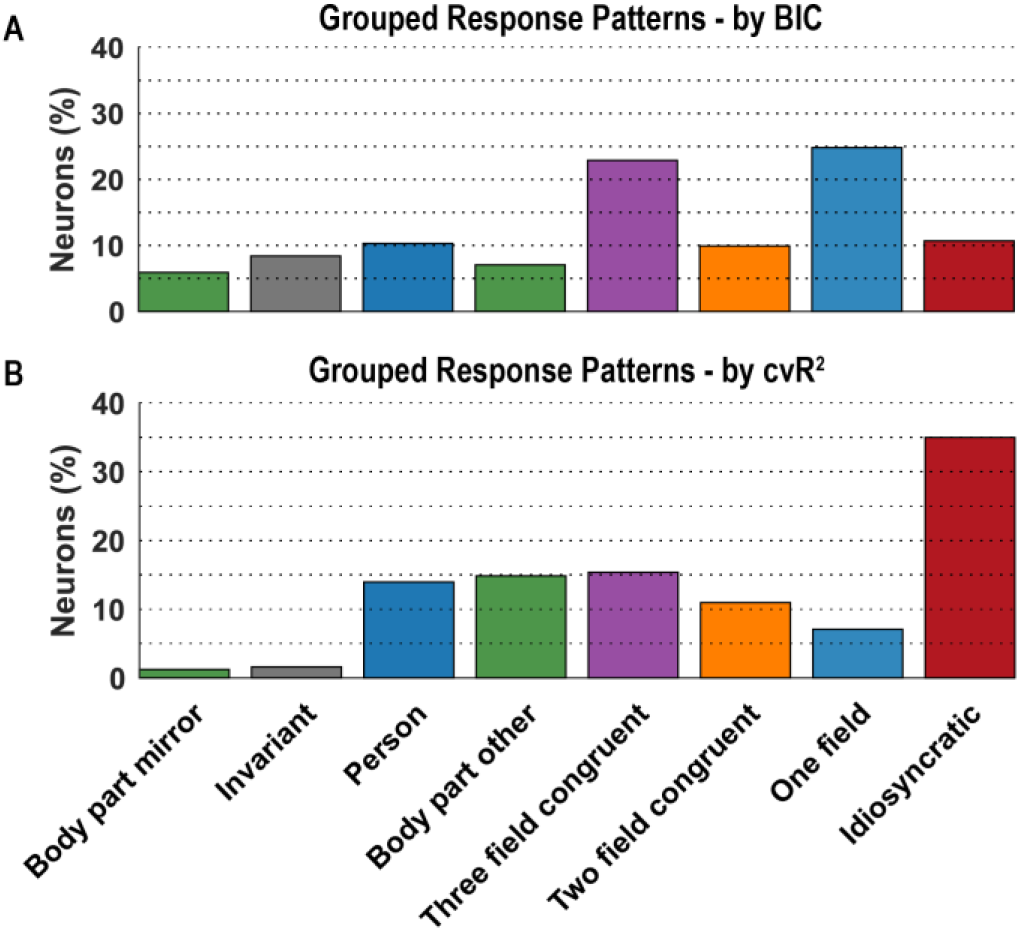
Summary of categorizing single-units into 51 possible patterns of correspondence across sensory fields based on BIC and *cvR*^2^. Related to figure 3. **A**, Histogram showing the percentage of PPC neurons that behaved according to each of the 8 categories of linear models (see Fig S3). Here the breakdown is shown based on using the Bayesian Information Criterion (BIC) as the metric for evaluating which of the 51 models best matched each neuron’s behavior. The category name is listed below each bar. **B**, Similar to A, except here the breakdown is based on the cross-validated coefficient of determination (*cvR*^2^).

## Movie Captions

**Movie S1: Example of a neuron demonstrating a mirror-like response.** This example neuron activated when NS felt a touch to her outer shoulder but did not activate when she felt a touch to her inner shoulder. This represents an example of the specificity of neural response. In addition, the neuron activated when NS visually observed a touch to the experimenter’s outer shoulder but did not activate when observed a touch to the experimenter’s inner shoulder. The fact that response properties of experienced and observed tactile sensations were similar demonstrates the congruency of the neural response.

**Movie S2: Example of a neuron demonstrating a mirror-like response.** This example neuron activated when NS felt a touch on her cheek and when she observed a touch on another person’s cheek, but not when she observed a touch on the cheek of a Styrofoam head.

